# Wide Window Acquisition and AI-based data analysis to reach deep proteome coverage for a wide sample range, including single cell proteomic inputs

**DOI:** 10.1101/2022.09.01.506203

**Authors:** Rupert L. Mayer, Manuel Matzinger, Anna Schmücker, Karel Stejskal, Gabriela Krššáková, Frédéric Berger, Karl Mechtler

**Affiliations:** Research Institute of Molecular Pathology (IMP), Vienna BioCenter, Vienna, Austria; Gregor Mendel Institute of Molecular Plant Biology (GMI), Austrian Academy of Sciences, Vienna BioCenter (VBC), Vienna, Austria; Institute of Molecular Biotechnology (IMBA), Austrian Academy of Sciences, Vienna BioCenter (VBC), Vienna, Austria

## Abstract

A comprehensive proteome map is essential to elucidate molecular pathways and protein functions. Although great improvements in sample preparation, instrumentation and data analysis already yielded impressive results, current studies suffer from a limited proteomic depth and dynamic range therefore lacking low abundant or highly hydrophobic proteins. Here, we combine and benchmark advanced micro pillar array columns (µPAC™) operated at nanoflow with Wide Window Acquisition (WWA) and the AI-based CHIMERYS™ search engine for data analysis to maximize chromatographic separation power, sensitivity and proteome coverage.

Our data shows that µPAC™ columns clearly outperform classical packed bed columns boosting peptide IDs by up to 140%. Already at classical narrow isolation widths CHIMERYS™ boosted ID rates by a factor of 2.6 compared to the conventional search engine MS Amanda 2.0. By combining CHIMERYS™ with WWA, even a 4.6-fold increase in ID rates could be achieved.

Using our optimized workflow, we were further able to identify more than 10,000 proteins from a single 2 h gradient shotgun analysis. We further investigated the applicability of WWA for single cell inputs and found that the choice of the optimal isolation window width depends on sample input and complexity. Using a short 5.5 cm column and very high flow rates during loading and column equilibration we improved sample throughput to ∼100 samples per day while maintaining high protein ID numbers. We believe that this is especially important for the single cell field where throughput is one of the most limiting factors.

Finally, we applied our optimized workflow on immunoprecipitations of Smarca5/SNF2H and found 32 additional interaction partners compared to the original workflow utilizing a packed bed column. These additional interaction partners include previously described interaction partners of Smarca5 like Baz2b as well as undescribed interactors including Arid1a, which is also involved in chromatin remodeling and has been described as key player in neurodevelopmental and malignant disorders.

## INTRODUCTION

Investigating the entire proteome, which defines the cell identity is one of the major objectives in the field of proteomics. A comprehensive proteome map is also important to elucidate entire molecular pathways. Improvements in sample preparation techniques and LC-MS instrumentation over the past decades enabled to identify thousands of proteins per sample. Powerful chromatographic separation to reduce sample complexity is therefore crucial to increase proteomic coverage. However, a high degree of separation is usually achieved by time-consuming long gradients, that create a bottleneck limiting throughput and sensitivity due to diluted signals based on broad peak widths.^1^ By applying offline fractionation, up to 12,000 proteins were successfully quantified in complex human samples.^2^ Fractionation is however a time-consuming step that might not be feasible for projects dealing with limited sample amounts or MS instrument time. Although high protein identification numbers are already reported from shotgun experiments from single cells^3^, those studies usually cover exclusively the most abundant proteins within the cell.^4^ Such a low resolution hinders relevant studies investigating specific post translational modifications (PTMs), transcription factors or other regulators present at a low copy number. Missing values across replicates are problematic for lower abundant peptide species as their respective precursor ions are only stochastically triggered in a DDA method.^5^ With increasing dataset size missing values become even more prevalent, which is why modern data analysis strategies try to alleviate this issue by matching precursor ions on the MS1 level across runs with the remaining empty values typically imputed by low numbers to allow statistical testing .^6,7^ This however comes with additional difficulties in FDR estimation and potentially mitigates proper quantitative accuracy. Another strategy reported to lower the fraction of missing values is data independent acquisition (DIA), acquiring all theoretical fragment ion spectra in sequential windows.^8^ The resulting data is however harder to analyze, as longer cycle times lead to less datapoints per peak to be used for label-free quantification. In combination with multiplexed samples using isobaric isobaric labels, the co-isolation of many precursor ions at once also impedes correct annotation of reporter ions to peptides and hence accurate quantification.

In this study we combine a multitude of approaches in order to overcome the above-mentioned bottlenecks. We benchmark advanced chromatographic setups that are based on microfabricated pillar array columns (µPAC™). In contrast to packed bed columns, these µPAC™ columns consist of highly ordered pillar arrays resulting in exceptionally homogenous flow paths, thus reducing peak broadening giving rise to sharper peaks and higher signal intensities.^9^ Furthermore, the use of superficially porous material on the surface of these micropillars instead of classical fully porous beads reduces the persistent adsorption of hydrophobic peptides, thereby reducing carry-over.^10^ By using rectangular shaped pillars, typical flow path lengths around 50 cm can be realized in very short physical column lengths of only 5.5 cm resulting in low backpressure at nano flow rates and a broad dynamic range of available flow rates ≤2.5µL/min. This enables fast loading, washing and conditioning of the column pre- and post-analysis increasing sample throughput.^10,11^

This is combined with a wide window acquisition (WWA) DDA method, which merges the strengths of DDA and DIA. As indicated by the name, we use a broad isolation window of ≥4 m/z in an otherwise data dependent precursor selection. In this way, precursor ions close to the selected ones are co-fragmented leading to chimeric spectra as would be recorded in a DIA run. This not only boosts ID numbers, but also improves coverage of low abundant peptides, which would otherwise be missed. This WWA approach particularly unfolds its potency when combined with the AI-driven search algorithm CHIMERYS™, which allows the confident identification of up to 11 peptides and more from a single chimeric spectrum. CHIMERYS™ was recently developed by MSAID and makes use of accurate fragment spectrum predictions trained on millions of spectra, which allows to utilize additional spectral properties such as relative signal intensities for drastically improved identification rates and accuracy.^12^

Implementation of all these innovations allowed us to reach unprecedented sensitivity, proteome coverage and improved throughput, which potentially reduces missing values, and allows novel biological insights.

## RESULTS

### Combination of micropillar array columns (µPAC™) with Wide Window Acquisition (WWA) and AI-driven data analysis results to reach unprecedented proteomic coverage

In our analytical platform, we combine recent technological innovations in liquid chromatography, mass spectrometry and data analysis to further improve the depth of analysis as illustrated in Figure 1A. We utilize the high versatility offered by the Vanquish™ Neo LC system, which offers a high flow rate range from a recommended 100nL/min up to 100µL/min without the need to install different flow meters. This is particularly helpful for columns that sustain high flow rates during sample loading or after analysis during washing and equilibration to reduce overhead times and improve sample throughput directly needed for the analysis of larger biological studies or clinical cohorts.

**Figure 1:**
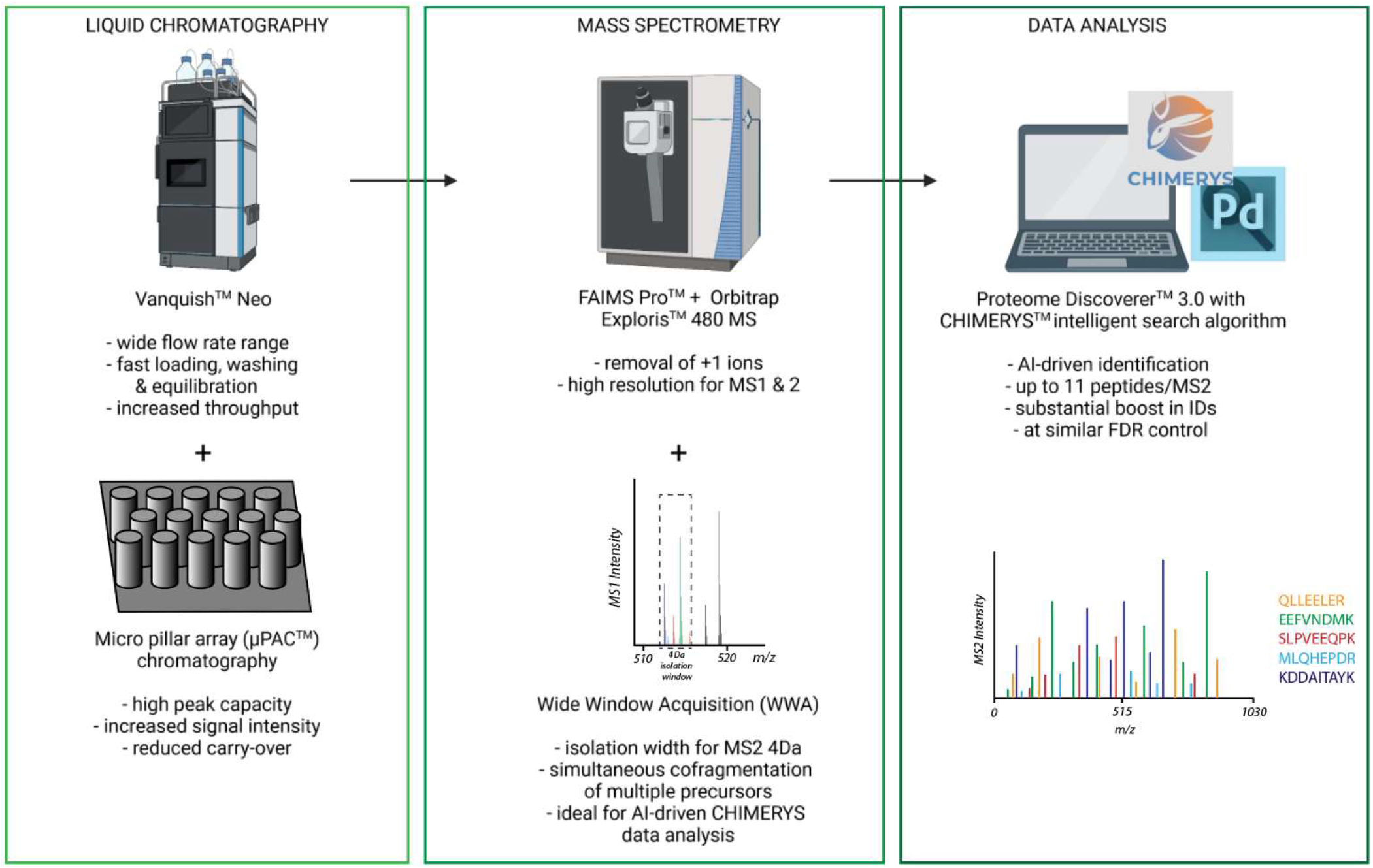
Combination of cutting-edge technological advancements in liquid chromatography, mass spectrometry and data analysis to achieve unprecedented proteomic depth. Scheme of the employed workflow including the use of the novel Vanquish™ Neo LC system and innovative µPAC™ columns, next to state-of-the-art FAIMS Pro™ and Orbitrap Exploris 480 mass spectrometry in conjunction with wide window acquisition (WWA), novel ProteomeDiscoverer™ 3.0 and the ground-breaking AI-driven CHIMERYS™ search algorithm. Created with BioRender.com.

The use of µPAC™ columns allows to reduce peak broadening due to minimized Eddy diffusion. These columns can cope with higher flow rates due to their unique design, which reduces backpressures as compared to packed bed columns, making them ideally suited for use on the Vanquish™ Neo LC system for fast loading, washing and equilibration at higher flow rates.

Mass spectrometric detection is aided by a FAIMS Pro interface, which acts as an ion filter based on charge state, molecular shape, conformation, and size to reduce the influx of undesired, singly charged background ions and thus improves signal to noise ratio of resulting spectra. In contrast to typical data dependent acquisition (DDA) approaches, which classically employ narrow precursor isolation windows in the range of 0.7-1.5 Da, we utilize WWA with precursor isolation windows of 4 Th and wider. We use these wide isolation windows as the CHIMERYS™ downstream data analysis tool can accurately and sensitively identify a high number of peptides from chimeric spectra. CHIMERYS™ typically surpasses the number of protein and peptide IDs detected by classical search engines substantially due to the inclusion of relative fragment intensities and other parameters often neglected in other search engines. CHIMERYS™ is embedded in Proteome Discoverer™ 3.0, which provides raw data pre-processing as well as further data analysis capabilities and statistical options.

This advanced analysis platform was utilized to measure a number of different samples including 12.5 ng HeLa digest with added standard peptides (Q4L), three different triple proteome mixes consisting of tryptic HeLa, yeast, and *E. coli* digests prepared in 0.1% TFA at ratios of 8:0.2:1.8 as well as 8:1:1 and 8:2:0. 400 ng of total peptide material were injected onto a 110 cm prototype 2^nd^ generation micropillar column installed on the Vanquish™ Neo applying a 120 min gradient.

### Micropillar columns (µPAC™) outperform packed bed columns

To assess column performance we used a QC standard mix from a HeLa digest including reference peptides in use within the Core for Life alliance^13^ to benchmark the 110 cm prototype 2nd generation micropillar column (GEN2) to commercial packed bed columns from Thermo (PepMap™, 50 cm bed length) and Waters (NanoEase, 25 cm bed length). As illustrated in Figure 2, the advanced GEN2 column outperforms its competitors by increasing protein and peptide IDs by ≤66 % and ≤140%, respectively. Based on these exciting numbers, we decided to further investigate and benchmark different µPAC™ columns, gradient lengths, and input amounts to get a comprehensive overview of column performance in a broad variety of sample conditions.

**Figure 2:**
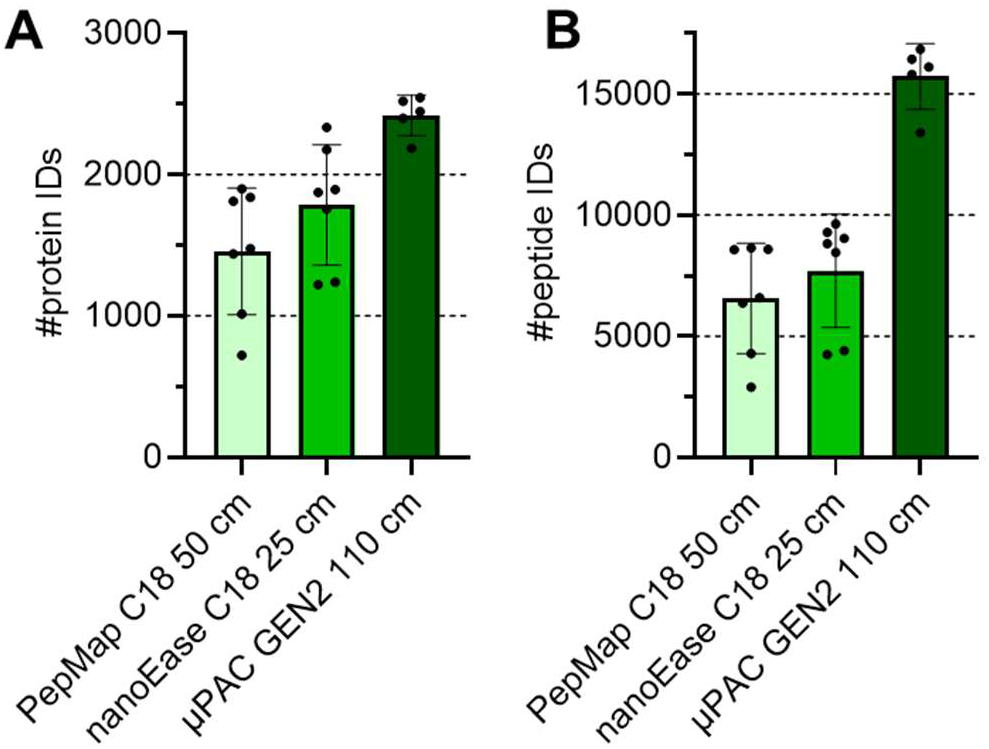
Benchmarking packed vs µ pillar columns. 12.5 ng of a HeLa QC mix were injected each using a trap and elute setting. Peptides were separated over 30 min using a linear gradient over 30 min from 1 – 35 % buffer B (80 % ACN, 0.01% TFA. Data acquisition using a standard DDA method with an isolation window of 1 Th and data analysis using CHIMERYS™ at 1% FDR on peptide and protein level, n = 6 technical replicates. **(A)** Protein and **(B)** peptide identifications are visualized.

### Different µPAC™ columns optimally facilitate a variety of proteomic applications and allow the identification of more than 10,000 proteins from a single run

To compare the performance and best suited application per column, three different µPAC™ columns were assessed including a 5.5 cm prototype column, a 50 cm Neo column as well as a 110 cm GEN2 column. Triple proteome mixes of tryptic HeLa, yeast, and *E. coli* digests were prepared at a ratio of 8:1:1 in 0.1% TFA at three different peptide concentrations 10 ng/µL, 100 ng/µL and 400 ng/µL. 0.5-1 µL of the lysate mix was injected directly into the analytical column without a trapping column using the Vanquish Neo LC system. Columns were grounded via the metal case of the LC system to prevent charging of the pillar array to avoid losses in separation power. Injection amounts were varied between 10 ng and 400 ng of total peptide material and gradient lengths were assessed up to 120 min. While the 5.5 cm prototype column and the 110 cm GEN2 column were connected to the Nanospray Flex™ PepSep sprayer via a custom-made connecting capillary (20 µm ID x 360 µm OD, length 20 cm), the 50 cm Neo column was connected directly to the sprayer via its nanoViper™ fittings.

Depending on the column length and the available flow rates, different gradient lengths per column type were tested including 5-60 min for the 5.5 cm prototype column, and 30-120 min for the 50 cm Neo and the 110 cm GEN2 column. Figure 3A visualizes the protein identifications obtained on average from these runs with a minimum of 1,934 protein IDs for the 5 min gradient on the 5.5 cm column and a maximum of 10,487 protein IDs for the 50 cm Neo column. For all measurements excellent reproducibility was achieved as indicated by the small error bars.

**Figure 3:**
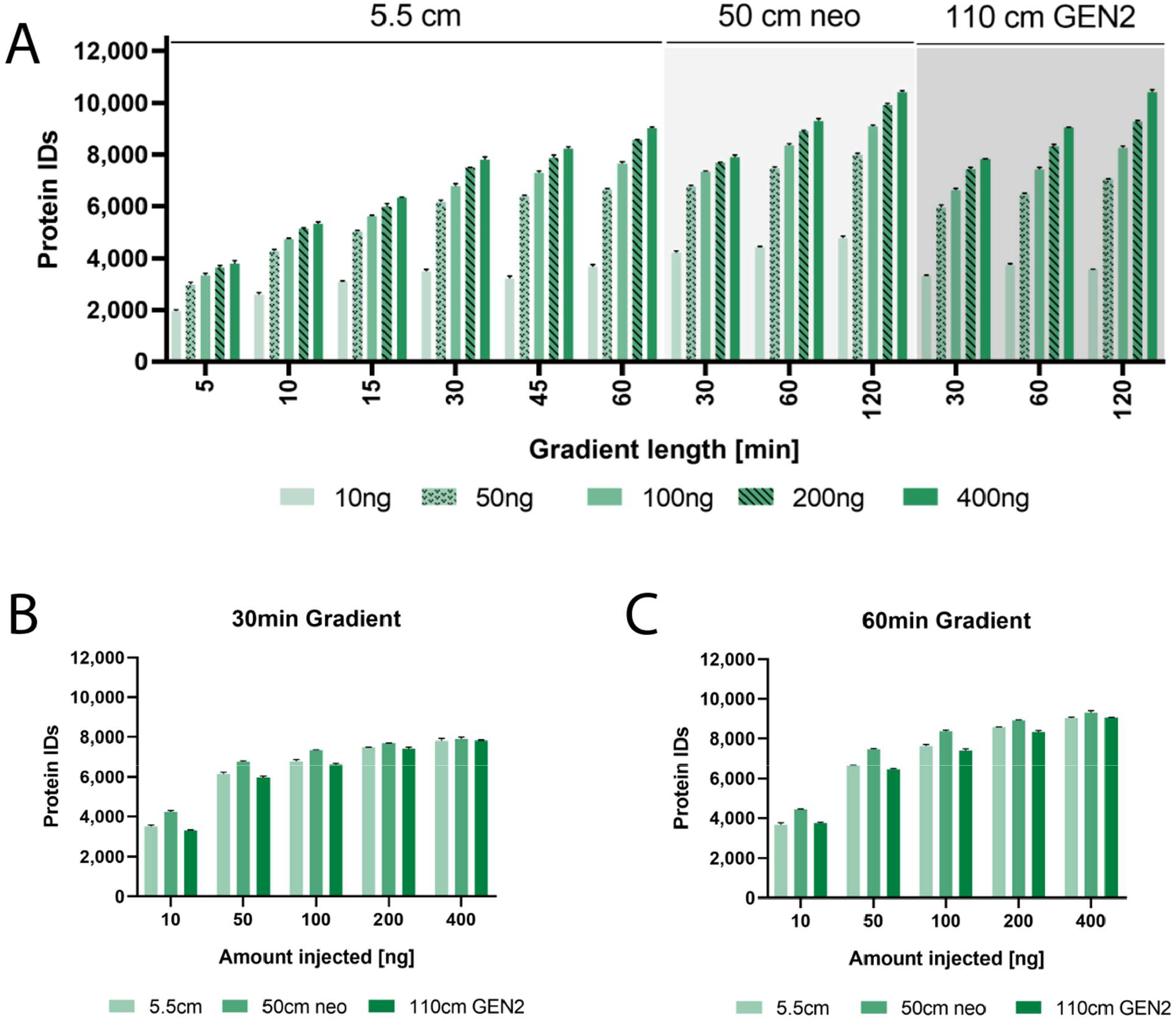
Benchmark of µPAC™ column designs and gradient lengths. (**A**) Bars indicate average number of identified proteins at 1 % FDR on protein level when acquiring using different column lengths, gradient times and input amounts indicated of a 8:1:1 H:Y:E proteome mix. Error bars indicate standard deviations, n = 2-3 technical replicates **(B)** Comparison of all columns for 30 min gradient from 10-800 ng peptide load. **(C)** Comparison of all columns for 60 min gradient from 10-800 ng peptide load. Data for panels B and C is already presented in panel A but displayed differently for easier visual comparison.

Unsurprisingly, the 120 min gradient with the two long columns achieved the greatest proteome depth with over 10,000 protein IDs on average for 400 ng injection amount. As expected, a general trend with higher injection amounts and longer gradient lengths towards higher proteome coverage was observed for all columns. To our surprise, a striking protein ID gap between an injection amount of 10 ng and 50 ng was observed, which had not been reported by earlier studies in our lab.^10^ In addition, increasing the peptide load led to a less pronounced increase in protein IDs. Only for longer gradients, the protein ID gap between different peptide loads showed a wider spread.

When comparing the columns for the 30 min and 60 min gradients, it became evident that the 50 cm Neo column performed best with the highest number of protein IDs. Particularly at 10 ng with the 30 min gradient it outperformed the 5.5 cm and 110 cm column with 4,222 over 3,506 and 3,316 protein IDs, respectively. At higher sample loads and longer gradients this gap was reduced, as shown for the 60 min gradient at 400 ng injection amount at 9,309 protein IDs for the 50 cm Neo column in contrast to 9,037 and 9,058 protein IDs for the 5.5 and 110 cm column, respectively. Eventually, for the 120 min gradient at 400 ng, there was no difference in protein IDs between the 50 cm Neo column and the 110 cm GEN2 column (10,421 protein IDs on average for both).

This increased sensitivity at low sample amounts but similar performance at higher sample loads for the 50 cm Neo column could be explained by the architecture of the column and the nanoviper fittings of the 50 cm Neo column (that was not equal the other two columns) allowing direct coupling to the valve and the pepsep sprayer with reduced post-column volume reducing peak broadening. As expected, this was particularly pivotal at low concentrations and became less important at high sample loads.

The 5.5 cm and 110 cm GEN2 columns perform similarly for the 30 and 60 min gradients. While the longer 110 cm GEN2 column intuitively seems to be better suited for higher sample loads and longer gradients, the 5.5 cm prototype column can also be readily used for very short runs utilizing gradient times of only 10 or even 5 min.

We therefore consider the 50 cm Neo column ideal as an all-round column that is ideally suited for lower as well as intermediate sample loads of 10-500 ng and shorter to intermediate gradient lengths around 15-120 min. The 5.5 cm prototype column in contrast seems to be best applied for shorter gradients due to the short physical length of the column and the high maximum flow rate up to 2.5 µL/min. The 110 cm GEN2 column did not reach its sweet spot in our tests so far but shows a tendency towards longer gradients of 120 min or longer and high sample loads of ≥400 ng.

### Unique column architecture of 5.5 cm protype column enables reduced run-to-run times resulting in superior sample throughput

While typically longer packed bed capillary columns have very limited maximum flow rates of up to ∼500 nL/min, the 5.5 cm prototype column offers a maximum flow of 2.5 µL/min. This column features rectangular-shaped micropillars in the form of bricks resulting in highly orthogonally prolonged flow paths and relatively large interpillar distances. This allows for highly efficient sample loading, washing, and conditioning in direct injection mode without the use of a trapping column. This substantially reduces the analysis overhead times and leads to more efficient use of the mass spectrometer as indicated in Figure 4A. The analysis of protein IDs/min run-to-run time for a 30 min gradient method revealed that, while the 50 cm Neo and the 110 cm GEN2 column could achieve a best of 108 and 86 protein IDs/min run-to-run time, respectively, the 5.5 cm prototype column delivered substantially more IDs/min at 147 representing a gain of 36% and 71% over the 50 and 110 cm column, respectively. When further cutting the gradient length down to 5 min, using the shortest column (Figure 4B) acquisition of up to 250 protein IDs/min and up to 96 sample injections per day are achievable.

**Figure 4:**
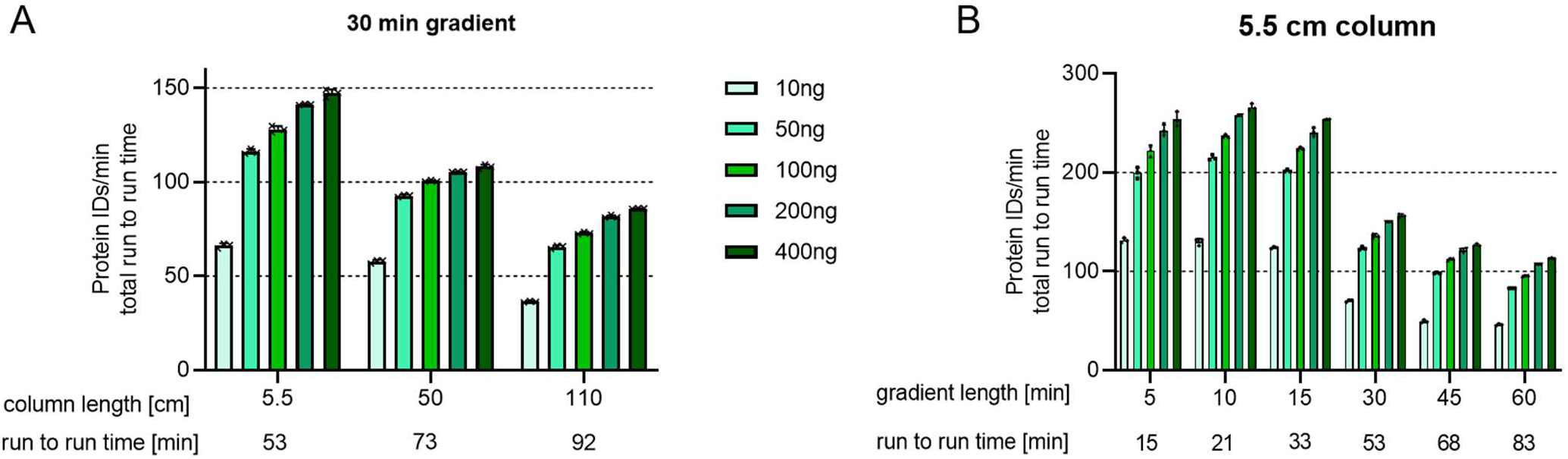
Assessment of protein IDs/min. Identified proteins at 1% protein FDR were normalized to the complete time between each injection (“run-to-run time”) to evaluate how efficiently the mass spectrometer was utilized over the entire duration of the run, .(**A**) Comparison of column types using a fixed gradeint length (**B**) Comparison of different gradient lengths using the 5.5 cm column.

Particularly for short or very short gradients of ≤30 min, reduced overhead times become more and more impactful as the overhead times remain stable while the total analysis time is reduced thereby consuming a great relative fraction of the total analysis time as illustrated in Table 1. At a flow rate during the analysis of 300 nL/min, the 110 cm GEN2 column offers little room for speeding up sample loading and equilibration as the maximum flow rate for this column is specified around 400 nL/min. The 5, 10 and 15 min gradient lengths were not measured on the 110 cm prototype column due to the low maximum flow rate. In contrast, the 5.5 cm prototype column allows flow rates up to 2,500 nL/min, offering considerably accelerated sample loading and column equilibration yielding up to 96 sample runs per day for very short 5 min gradients.

**Table 1:** Different column lengths and geometries result in substantially variable overhead times. All gradients used for the 5.5 cm prototype column are presented with optimized overhead times in comparison to standard overhead times for the 110 cm GEN2 column. Overhead times include sample pickup and loading as well as equilibration after washing. Due to the high maximum flow rate for the 5.5 cm column, overhead times can be dramatically faster allowing sample throughputs of up to 96 samples per day. Of note, 5, 10 and 15 min gradients were not realized using the 110 cm column due to its limited flow rate and the 120 min gradient was not realized on the 5.5 cm column.

| Gradient length [min] | Wash time [min] | Total over-head [min] | Run-to-run time [min] | Samples per day no washes or QCs | % over-head of run-to-run time | Total over-head [min] | Run-to-run time [min] | Samples per day no washes or QCs | % over-head of run-to-run time |
| --- | --- | --- | --- | --- | --- | --- | --- | --- | --- |
| all columns |  | 5.5 cm column |  |  |  | 110 cm GEN2 column |  |  |  |
| 5 | 4 | 6 | 15 | 96 | 40 | 47 | - | - | - |
| 10 | 4 | 7 | 21 | 69 | 33 |  | - | - | - |
| 15 | 10 | 8 | 33 | 44 | 24 |  | - | - | - |
| 30 | 15 | 8 | 53 | 27 | 15 |  | 92 | 16 | 51 |
| 45 | 15 | 8 | 68 | 21 | 12 |  | 107 | 13 | 44 |
| 60 | 15 | 8 | 83 | 17 | 10 |  | 122 | 12 | 39 |
| 120 | 15 | - | - | - | - |  | 182 | 8 | 26 |

### Utilization of AI-driven search engine CHIMERYS™ for improved proteomic depth over a state-of-the-art search engine

For better utilization of all information provided within fragment spectra, we evaluated the use of the AI-driven, groundbreaking search engine CHIMERYS™. Next to the m/z values of peptide fragments, CHIMERYS™ can harness additional spectral information such as relative abundance of fragment signal, which allows the accurate identification of numerous peptides from highly chimeric fragmentation spectra resulting from our proteome-mix representing a highly complex sample. **Figure 4A** clearly highlights that CHIMERYS™ offers improved proteomic depth compared to a more classical state-of-the-art search engine with an additional 68% peptide IDs and 32% protein IDs for a typical 1 Th isolation window. Even more pronounced was the improvement on the identification rate assigning 2.6 times as many PSMs to MS/MS spectra in comparison to MS Amanda 2.0. ^14,15^ Using a wider precursor isolation window of 4 m/z, which we relate to as wide window acquisition (WWA), we generated richer, more complex MS2 spectra, further favoring the advantage of CHIMERYS™ to decipher highly chimeric fragmentation spectra. This boosted the identification rate of CHIMERYS™ to around 125% meaning that on average there was more than one PSM per MS/MS spectrum identified, corresponding to a 4.6-fold boost over the classical search engine employed, while peptide IDs were boosted by 113% and protein IDs by 41%. Figure 4B indicates that these additional protein IDs substantially overlap with the results from MS Amanda 2.0, thereby conserving confident IDs, while detecting low abundance proteins.

### The ideal precursor window size to optimally utilize CHIMERYS™ differs according to sample abundance

We tested an array of different precursor isolation widths as illustrated in Figure 6 for different input amounts of tryptic HeLa digest from 250 pg up to 400 ng. Deducing from these data, we could assign 4 m/z as ideal isolation width for maximum protein IDs for standard injection amounts (200-400 ng), while the best isolation width for low input samples was around 8 - 12 m/z. Considering the reduced complexity that is to be expected from low input samples, it seems intuitive that wider isolation windows are beneficial to reach similar levels of complexity in the fragmentation spectra. Indeed, our data supports the advantage of broader isolation windows for lowered input amounts (see Figure 6C). This trend is even more pronounced on the peptide level or when investigating the number of identified peptides per spectrum (identification rate) (Supplemental Figure 1). Hence, for best utilization of the CHIMERYS™ search engine it appears beneficial to optimize the precursor isolation window according to sample complexity and injection amount to obtain optimal results.

**Figure 5:**
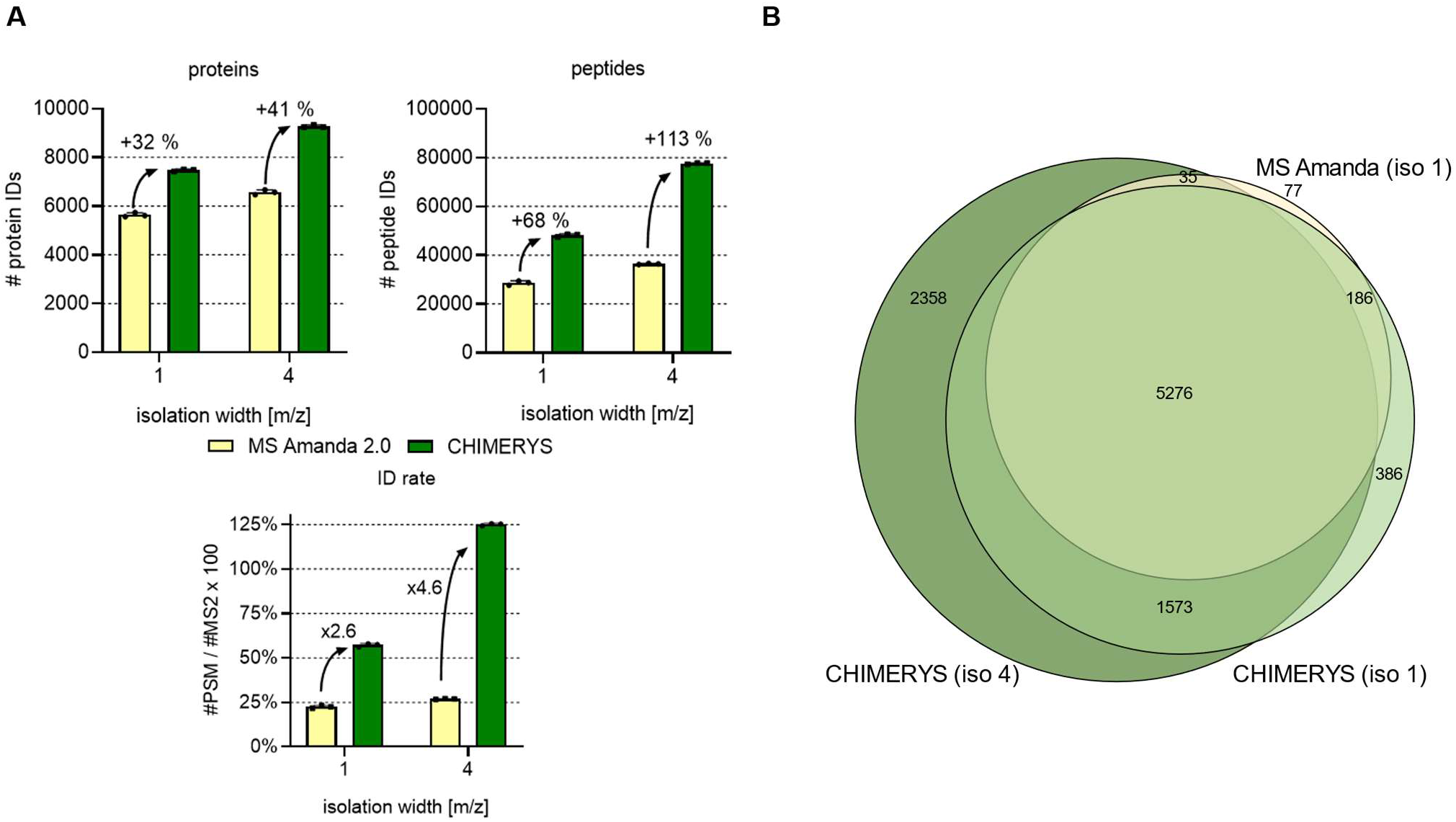
AI-driven search engine CHIMERYS™ improves peptide ID rates substantially to boost protein IDs. 200 ng proteome-mix H:Y:E = 8:1:1 were separated over 120 min using the 110 cm GEN2 column prior to MS acquisition and data analysis with the indicated strategy, n =3 technical replicates **(A)** Typical DDA measurements show improved ID rates on peptide as well as protein level using CHIMERYS™, this boost is even more pronounced when using WWA that uses precursor isolation widths of 4 m/z. **(B)** CHIMERYS™ identifies the same proteins as well as further species extending the accessible proteomic coverage of the sample under analysis.

**Figure 6:**
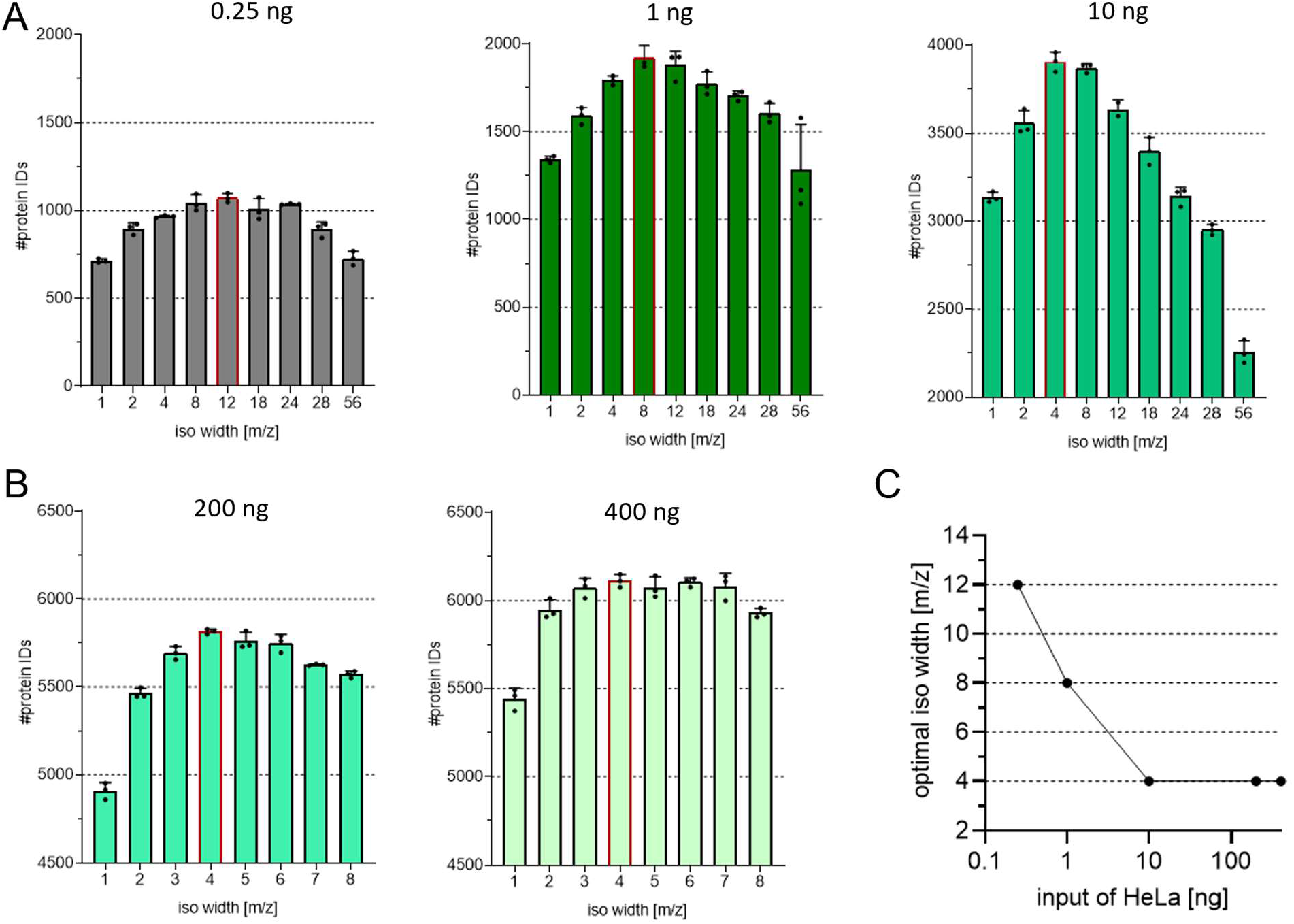
Ideal precursor window size for wide window acquisition to optimally utilize CHIMERYS™ depends on sample abundance. Different isolation window sizes were tested to identify the most well-suited precursor isolation window size for **(A)** low sample input such as 250 pg up to 10 ng measured on the 5.5 cm column and **(B)** standard sample input from 200 to 400 ng leading measured on the 50 cm column. (**C**) Optimal isolation width in dependence of used input amount.

### CHIMERYS™ allows to access a wider dynamic range of tryptic peptides for a more comprehensive proteomic picture

We speculated that using CHIMERYS™ and WWA, we could identify peptides with a broader dynamic concentration range, which would be the reason for the increased detection depth. To this end, PSMs were ranked according to intensity and assigned to a PSM index from the lowest abundant PSM starting at PSM index 1 up to the highest abundant PSM with the highest PSM index as displayed in **Figure 6**. This plot visualizes that the combination of CHIMERYS™ and WWA unlocks a broader dynamic range of peptides for identification, which in turn allows the identification of lower abundant proteins.

### Investigation of AP-MS samples with the optimized analysis platform unlocks additional biological insights

To address the impact of our optimized platform on the investigation of protein-protein interactions, we analyzed six mouse co-immunoprecipitation samples using the µPAC™ 110 cm GEN2 column connected to an Orbitrap Exploris 480 interfaced with a FAIMS Pro device using 4 Th WWA, Proteome Discoverer 3.0, and CHIMERYS™ for data analysis. The co-immunoprecipitation samples were generated using anti-flag M2 beads specifically enriching for the flag-tagged bait protein. Three samples had been generated using the flag-tagged bait protein, while the other three included non-tagged, wild type bait protein, serving as the control group. The objective of the analysis was the identification of specific interactors to the Smarca5/Snf2h protein in mouse, which is known to be part of numerous complexes involved in chromatin-remodeling such as the WICH complex or the NoRC-5 ISWI chromatin remodeling com-plex.^16–18^

In good agreement with the elevated numbers of proteins identified we had seen during our initial tests, also substantially more protein IDs could be achieved for the co-immunoprecipitation samples. While a standard workflow with a 50 cm packed bed column, a 1 Th precursor isolation width and MS Amanda 2.0 resulted in 901 protein and 5,141 peptide IDs, our advanced platform enabled the identification of 2,175 proteins and 14,102 peptides as illustrated in Figure 8A and B. The use of the 110 cm GEN2 column allowed the detection of 59% more proteins and 72% more peptides confirming the high resolving power of µPAC™ and its applicability in biological projects. Interestingly, the improved chromatography allows CHIMERYS™ already at an isolation width of 1 Th to boost peptide and protein IDs more strongly at 26% more peptides and 21% more proteins when compared to the improvements reached with CHIMERYS™ for the packed bed column runs with only +4 and +5% for protein and peptide IDs, respectively. As shown earlier in Figure 4A and 5, CHIMERYS™ reaches its full potential only at higher isolation widths with an ideal isolation width of 4 Th for regular abundance samples, which again provides an over proportional boost in IDs of +61% protein and +76% peptide IDs, when using both µPAC™ and an isolation width of 4 Th over 1 Th.

**Figure 7:**
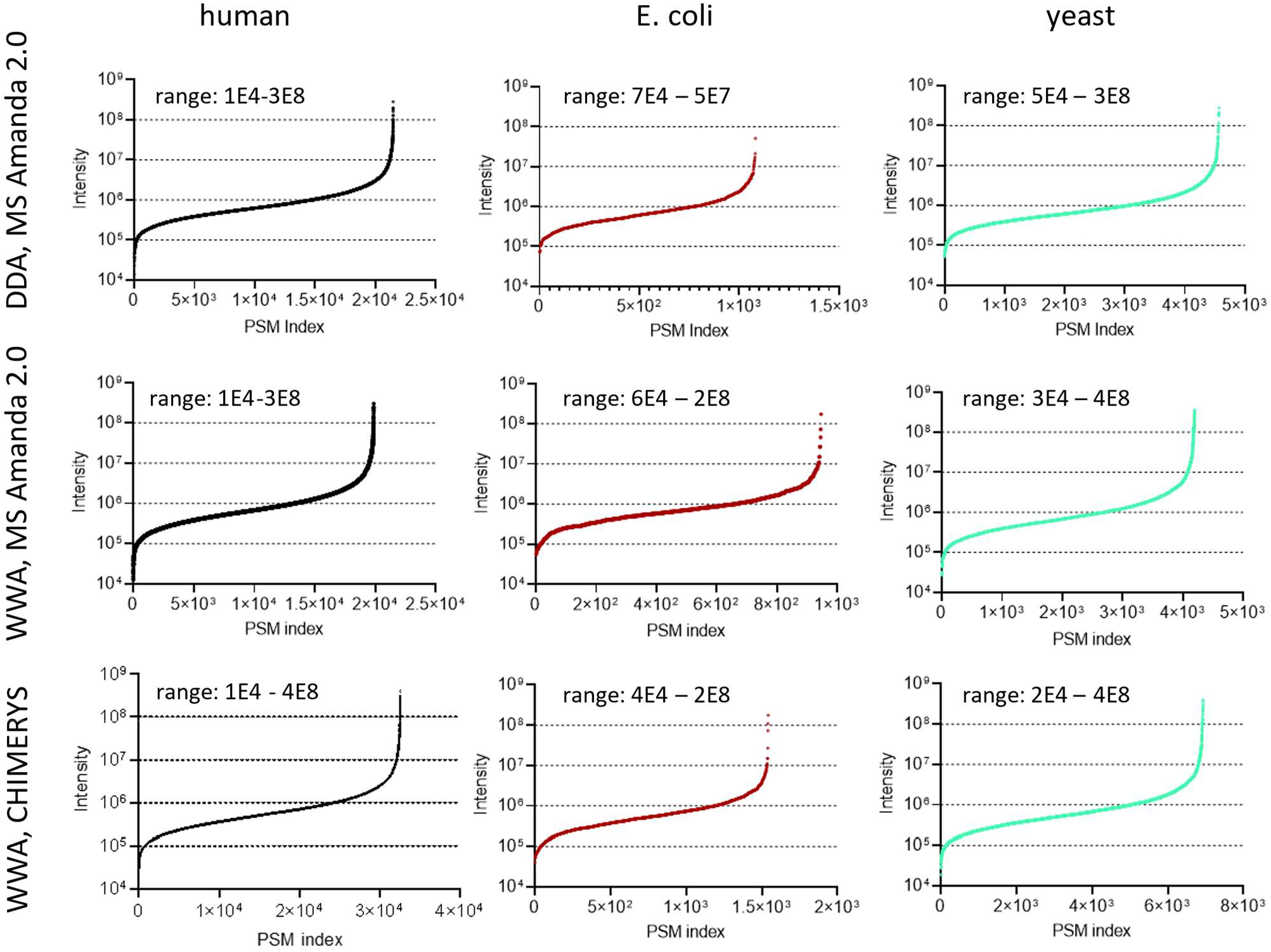
Data analysis using CHIMERYS™ improves the dynamic range of identifiable tryptic peptides. The identifiable peptide pools in the triple proteome mix samples between classical DDA acquisition using MS Amanda 2.0 and WWA using CHIMERYS™ were compared, indicating that the later approach allows the identification of peptides of a greater dynamic range in general when compared to the classical approach.

**Figure 8:**
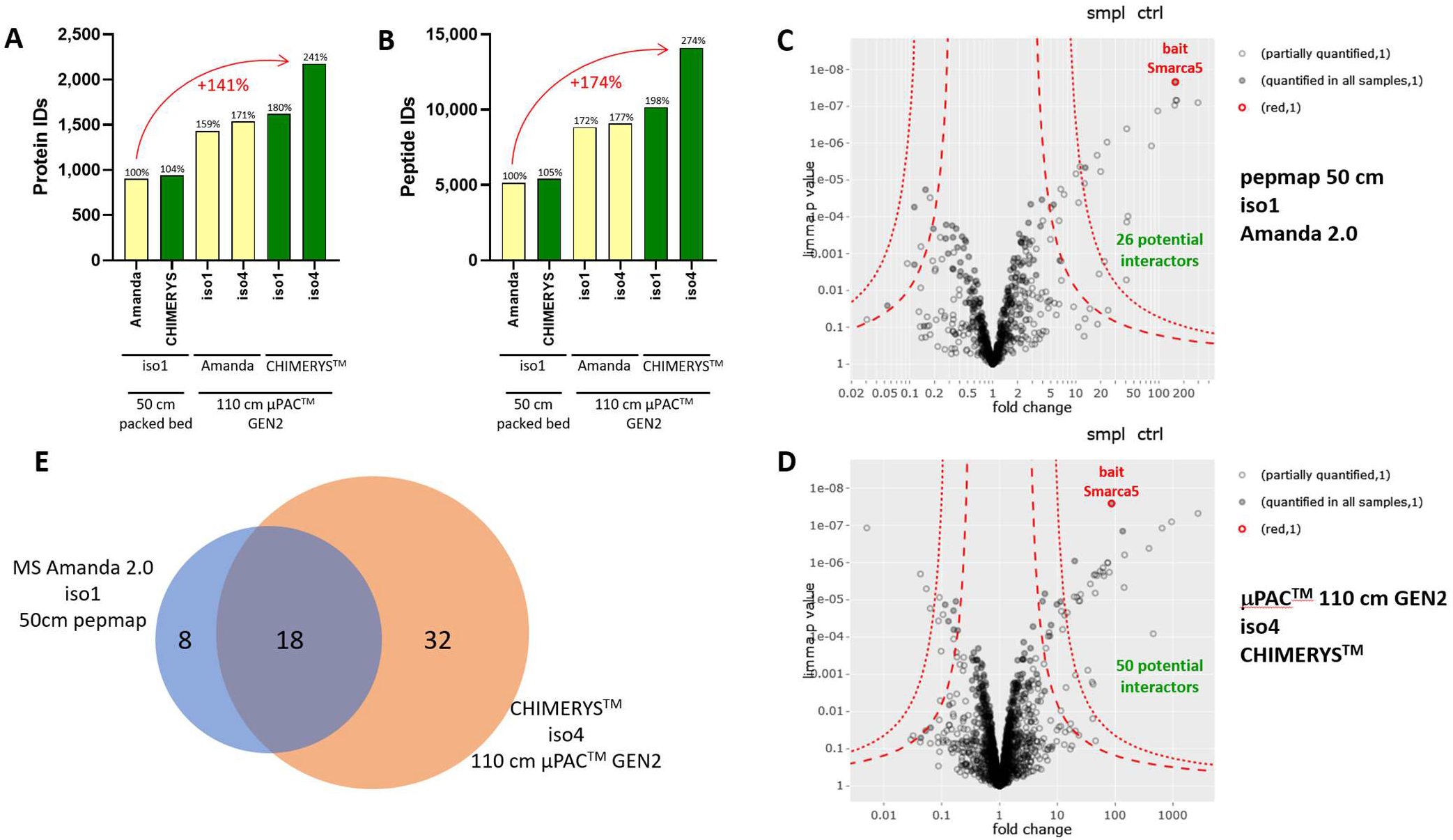
Implementation of micropillar array chromatography, wide window acquisition (WWA) and CHIMERYS™ facilitates deeper insights into protein-protein interactions. Six mouse co-immunoprecipitation samples were measured with three different conditions using i) a 50 cm PepMap™ column and 1 Th precursor isolation width, ii) a µPAC™ 110 cm GEN2 column and 1 Th precursor isolation width, and iii) a µPAC™ 110 cm GEN2 column with 4 Th precursor isolation width. **(A)** and **(B)** visualizes the gradual improvements on the analytical platform substantially increasing the number of identified proteins and peptides, respectively. By implementation of micropillar array chromatography in combination with WWA and CHIMERYS™ the number of protein and peptide IDs could be increased by 141% and 174%, respectively. **(C)** and **(D)** displays Volcano plots of the basic analytical approach and the advanced analytic platform, respectively, based on limma^19^ analyses to determine differential enrichment between flag-tagged and WT Smarca5 bait protein. The finely dashed lines indicate 1% FDR with the second line below indicating 5% FDR. The improvement in protein identifications is reflected in the number of detected potential interactors with 50 potential interactors for the advanced analytical platform versus 26 for the basic analytical approach. **(E)** The advanced analytical platform shows high overlap with the classical, basic approach, while identifying substantially more unique potential interactors.

These additional protein and peptide IDs of the optimized system in turn also allows the identification of substantially more potential interactions as indicated in Figure 8C and D. Using the statistical package LIMMA^19^, 26 and 50 potential interactors could be identified for the conventional and the advanced system, respectively, nearly doubling the number of potential interactors. Figure 8E illustrates that the majority of the interactors found by the conventional system could also be reproduced by our advanced platform, of which many are known binding partners of Smarca5 such as Rsf1^20^, Bptf^21^, Baz2a^16^, Cecr2^21^, Baz1b^21^ and Rbbp4.^22^ The eight proteins missed by the advanced platform contained five ribosomal proteins, which are frequently occurring, potential contaminants in affinity purification mass spectrometry experiments^23^ and did not feature any known Smarca5 interactors. In contrast, the additional advanced system-identified 32 interactors contained Baz2b, a Smarca5 binding partner within the BRF-5 ISWI chromatin-complex, as well as other proteins indicated in the STRING database to show interaction with Smarca5 such as Smarce1, Actl6a, Bud31 and Nanog.^24^ In addition, several other potential interactors that were identified show localization to the nucleus as expected when associated with the chromatin remodeling protein Smarca5 such as Dpf2, Znf280d, Nup62 as well as others, potentially hinting towards a high number of genuine Smarca5 interactors amongst these potential interactor candidates exclusively identified by our advanced analytical platform. Arid1a (BAF250a) was also exclusively detected as potential interactor in the advanced platform and has previously been described as component of the SWI/SNF chromatin remodeling complex. ARID1a (BAF250a) is a principal component of SWI/SNF (SWItch/Sucrose Non-Fermentable) family of evolutionary conserved, multi-subunit chromatin remodeling complexes. It interacts with other chromatin remodelers, but to date its interaction with Smarca5/ISWI had not been reported. Arid1a holds great clinical relevance as it has been associated with neurodevelopmental^25^ and malignant disorders^26,27^.

## CONCLUSION

In spite of many improvements of mass spectrometry-based proteomics in the last two decades, comprehensive sample throughput and analysis depth are still challenging. The comprehensive analysis of complex proteomic samples up to a depth of 12,000 proteins has been demonstrated by others. However, time consuming offline fractionation was required to achieve this level of comprehensiveness severely hampering sample throughput, which is typically required for large biological or clinical studies and is also of paramount interest for single cell proteomic measurements. In the current manuscript we describe an advanced workflow to routinely identify more than 10,000 proteins from a highly complex sample with a 120 min gradient. In addition, this workflow is also capable of delivering outstanding throughput of up to 96 samples per day when using the 5.5 cm prototype column in combination with short 5 min gradients offering reasonably good proteomic coverage of close to 4,000 protein IDs. We propose an optimized analysis platform combining cutting edge technologies including the Vanquish Neo LC system with micropillar array chromatography, the innovative WWA measurement strategy and the AI-based search engine CHIMERYS™ in conjunction with the recently launched Proteome Discoverer 3.0. This platform offers high flexibility for the comprehensive deep analysis of complex samples, or for the highly efficient high throughput analysis of large studies with hundreds of samples in several days. Input amounts of 250 pg reflecting single HeLa cells were also assessed displaying good proteomic coverage of over 1,000 protein IDs and benefitted from WWA in conjunction with CHIMERYS™. We also demonstrate how this advanced platform facilitates the analysis of protein-protein interactions by improving the detection of potential Smarca5 interactors by 92% from 28 to 50 as compared to a standard workflow. These novel potential interaction partners include many known Smarca5 interactors as well as previously unreported ones with clinical relevance such as Arid1a, that has been associated with neurodevelopmental as well as malignant disorders.

## METHODS

### Column benchmarking and WWA optimization studies

#### Sample preparation

##### QC mix for initial benchmarking with packed bed columns

LC-MS/MS Peptide Reference Mix (V7491, Promega) in HeLa digest (Thermo Scientific, Pierce™ HeLa Protein Digest Standard, 88328)

##### Triple proteome mix HYE

HeLa (H) (Thermo Scientific, Pierce™ HeLa Protein Digest Standard, 88328), yeast (Y) (Promega, MS Compatible Yeast Protein Extract, Digest, Saccharomyces cerevisiae, 100ug, V7461) and E. coli digests (E) (Waters, MassPREP E. coli Digest Standard, 186003196) were combined at a ratio of H :Y : E = 8:1:1, in 0.1% TFA.

##### HeLa samples

HeLa (Thermo Scientific, Pierce™ HeLa Protein Digest Standard, 88328) was diluted using 0.1% TFA to reach concentrations of 1ng/uL for 1 ng injections, 10 ng/uL for 10 ng injections, 200 ng/uL for 100 and 200 ng injections and 800 ng/uL for 400 and 800 ng injections respectively. To mimic single cell level injections HeLa digest was diluted to 250 pg/uL in 0.1% TFA including 5% DMSO and 1uL of this mix was used for injection.

Samples were prepared in glass autosampler vials (Fisherbrand™ 9mm Short Thread TPX Vial with integrated Glass Micro-Insert; Cat. No. 11515924). All liquid handling was done as fast as possible without unnecessary time gaps aiming to minimize sample adsorption on any surfaces.

#### Liquid chromatography-mass spectrometry (LC-MS) analysis

All samples were analyzed using a Vanquish Neo UHPLC operated in direct injection mode and coupled to the Orbitrap Exploris 480 mass spectrometer equipped with a FAIMS Pro interface (ThermoFisher Scientific). Analyte separation was performed using either the new generation prototype 110cm, 50 cm pillar array column or a 5.5 cm brick shape pillar column prototype (Thermo Fisher). For benchmarking, classical packed bed columns were used: nanoEase M/Z Peptide CSH C18 Column (130Å, 1.7 μm, 75 μm × 250 mm, Waters, Germany) or PepMap™ C18 (500 mm × 75 μm ID, 2 μm, 100 Å, Thermo Fisher Scientific). All columns were operated at 50°C and connected to an EASY-Spray™ bullet emitter (10 µm ID, ES993; Thermo Fisher Scientific) An electrospray voltage of 2.4 kV was applied at the integrated liquid junction of EASY-Spray™ emitter. To avoid electric current from affecting the upstream separation column, a stainless steel 50 µm internal bore reducing union (VICI; C360RU.5S62) was electrically connected to the grounding pin at the pump module.

Peptides were separated using gradients ranging from 5 min to 120 min ramping time as detailed in Supplementary Table 1

#### MS Acquisition

MS acquisitions was performed in data-dependent mode, using a full scan with *m/z* range 380-1200, orbitrap resolution of 60.000, target value 100 %, and maximum injection time set to auto. 1 to 4 FAIMS compensation voltages were combined in a single run as detailed in Supplementary Table 2 using a total cycle time of 3 sec. The intensity threshold for precursor was set to 1e4. Dynamic exclusion duration was based on the length of the LC gradient and is detailed in Supplementary Table 2.

Fragmentation by HCD was done using a normalized collision energy of 30 % and MS-Ms spectra were acquired at a resolution of 15000. Precursors were isolated in a window of 4 Th for WWA and 1 Th for normal DDA respectively and if not given other.

#### Data analysis

MS/MS spectra from raw data were imported to Proteome Discoverer (PD) (version 3.0.0.757, Thermo Scientific). Database search was performed using MS Amanda^24^ (version 2.5.0.16129) or CHIMERYS™ as indicated against a combined database of human (uniprot reference, version 2022-03-04, 20,509 entries), yeast (uniprot reference, version 2015-01-13, 4,877 entries) and E. coli (uniprot reference, version 2021-11-19, 4,350 entries) as well as common contaminants (PD_Contaminants_IGGs_v17_tagsremoved, 344 entries). For HeLa samples, yeast and *E. coli* databases were removed for searches. Trypsin was specified as proteolytic enzyme, cleaving after lysine (K) and arginine (R) except when followed by proline (P) and up to two missed cleavages were allowed. Fragment mass tolerance was limited to 20 ppm and carbamidomethylation of cysteine (C) was set as a fixed modification and oxidation of methionine (M) as a variable modification. Identified spectra were rescored using Percolator ^25^ and results were filtered for 1% FDR on peptide and protein level. Abundance of identified peptides was determined by label-free quantification (LFQ) using IMP-apQuant without match beween runs (MBR).^28^

### Analysis of immunoprecipitation samples

#### Generation of flag tagged Smarca5 cell line

For endogenous tagging of SMARCA5, WT cells grown on a 10 cm plate were transfected with sgRNA/Cas9 ribonucleoprotein complex and 15 µg of plasmid carrying a GFP-3XFLAG tag sequence flanked on both sides by 500 bp of Smarca5 stop codon adjacent sequence. For sgRNA/Cas9 ribonucleoprotein complex, sgRNA for Smarca5 was incubated with Cas9 in cleavage buffer for 5 min at RT. To transfect mouse ES cells, electroporation was carried out following the instructions of the Mouse Embryonic Stem Cell Nucleofector Kit from Lonza. After 2 days recovery, GFP expressing cells were FACS sorted and seeded for clone picking on 15 cm plate. The clones were individually picked and further grown. The resulting cultures were screened by FACS analysis.

#### Immunoprecipitation

Mouse ES cells were grown on 15 cm plates until confluency. Cells were harvested and washed with 1 × PBS. Then, cells were resuspended in buffer1 (10 mM Tris-HCl pH 7.5, 2 mM MgCl_2_, 3 mM CaCl_2_, Protease inhibitors (Roche)) and incubated for 20 min at 4°C. After centrifugation, cells were resuspended in buffer2 (10 mM Tris-HCl pH 7.5, 2 mM MgCl_2_, 3 mM CaCl_2_, Protease inhibitors (Roche), 0.5 % IGEPAL CA-630, 10% glycerol) and incubated for 10 min at 4°C. After this, cells were again centrifuged and nuclei were resuspended in buffer3 (50 mM HEPES-KOH pH 7.3, 200 mM KCl, 3.2 mM MgCl_2_, 0,25% Triton, 0.25% NP-40, 0.1% Na-deoxycholate, 1 mM DDT, Protease inhibitors (Roche)). 4 µl benzonase was added to the nuclei and the digest was incubated for 1 hour at 4°C. The lysate was cleared by centrifugation. For the Smarca5 IP, WT and Smarca5-Flag cells were added to magnetic anti FLAG M2 beads and incubated at 4°C for 2 hours. Beads were subsequently washed four times with buffer4 (50 mM HEPES-KOH pH 7.3, 200 mM KCl, 3.2 mM MgCl_2_, 0,25% Triton, 0.25% NP-40, 0.1% Na-deoxycholate, 1 mM DDT) and four times with Tris buffer (20 mM Tris-HCl pH 7.5, 137 mM NaCl).

#### On bead digest

Frozen magnetic beads were thawed, 20 μL of 100 mM ammonium bicarbonate (Merck, Sigma-Aldrich, 09830-1KG) as well as 600 ng of LysC (Wako Chemicals, 129-02541) added and incubated for 4h at 37°C at 1,200 rpm shaking. Supernatant was aspirated, transferred and tris-(2-carboxyethyl)-phosphin (TCEP, Merck, Sigma-Aldrich, 646547-10×1ML) added up to 1 mM and cysteine reduction performed for 30 min at 60°C. Reversible blockage of cysteines was performed with S-methyl methanethiosulfonate (MMTS, Merck, Sigma-Aldrich, 64306-1ML) at 4 mM for 30 min at room temperature. Trypsin digestion was performed overnight with 600 ng trypsin (Promega, V5280) at 37°C without shaking. Digestion was quenched by addition of 10 μL 10% trifluoroacetic acid (TFA, Thermo Scientific, VC296817).

#### LC-MS analysis of immunoprecipitation samples

The nano HPLC system used was an UltiMate 3000 RSLC nano system coupled to a Orbitrap Exploris 480 mass spectrometer, equipped with an EASY-spray ion source (Thermo Fisher Scientific) and a Jail-Break 1.0 adaptor insert as the spray emitter (Phoenix S&T) as well as a FAIMS Pro device (Thermo Scientific). Peptides were loaded on a trapping column (Thermo Fisher Scientific, PepMap™ C18, 5 mm × 300 μm i.d., 5 μm particles, 100 Å pore size) at a flow rate of 25 μl/min using 0.1% TFA as the mobile phase. 10 min after sample injection, the trapping column was switched in line with the analytical column and peptides were eluted from the trapping column onto the analytical column using a flow rate of 230 nl/min and a binary 3 h gradient of 220 min. The analytical column was either a classical 50 cm packed bed column (Thermo Fisher Scientific, PepMap™ C18, 500 mm × 75 μm i.d., 2 μm, 100 Å) or a prototype 110 cm µPAC™ GEN2 column (Thermo Fisher Scientific, micropillar array column, C18) The gradient started with mobile phases of 98% A (water:formic acid, 99.9:0.1 v/v) and 2% B (water:acetonitrile:formic acid, 19.92:80:0.08 v/v/v), increasing to 35% B over the next 120 min, followed by a gradient over 5 min to 95% B, held for 5 min and decreasing over 2 min back to gradient 98% A and 2% B for equilibration at 30°C. The trapping column was switched out of line from the analytical column 3 min after reaching 2% B again, and equilibration at 2% B was continued until the total run time of 165 min was reached.

The Orbitrap Exploris 480 mass spectrometer was operated in the data-dependent mode with the FAIMS Pro using three different compensation voltages (CVs) at -45, -60 and -75 in an alternating fashion switching between CVs every 0.9s. A full scan (m/z range of 350–1,200, MS1 resolution of 60,000, normalized AGC Target of 100%) was followed by MS/MS scans of the most abundant ions until the cycle time of 0.9s was reached. MS/MS spectra were acquired using a normalized collision energy of 30%, an isolation width of 1.0 *m/z*, a resolution of 30,000 and a normalized AGC target of 200%. Precursor ions selected for fragmentation (exclude charge state 1, 7, 8 and >8) were placed on a dynamic exclusion list for 45 s. Additionally, the intensity threshold was set to a minimum intensity of 2.5 × 10^4^.

### Data analysis

For peptide identification, RAW files were loaded into Proteome Discoverer (v.3.0.0.757, Thermo Scientific). All the created MS/MS spectra were searched either using MSAmanda v.2.0^14,15^ or CHIMERYS™ (MSAID GmbH, Germany). For the processing step, the RAW files were searched against the mouse Uniprot reference database (2022-03-04; 21,962 sequences and 11,728,099 residues) and a in-house contaminant database (PD-Contaminants_IGGs_v17_tagsremoved; 344 sequences and 142,046 residues).

The following search parameters were used for MS Amanda 2.0: the peptide mass tolerance was set to ±5 ppm and the fragment mass tolerance to 10 ppm; the maximal number of missed cleavages was set to 2; and the result was filtered to 1% false discovery rate (FDR) on the protein level using the Percolator algorithm integrated in Thermo Proteome Discoverer. Beta-methylthiolation on cysteines was set as fixed modification, whereas methionine oxidation was set as variable modification.

For CHIMERYS™ the following search parameters were used: as prediction model inferys_2.1_fragmentation was chosen using trypsin with a maximum of two missed cleavages. Peptide length was restricted to 7-30 amino acids, a maximum of three modifications per peptide and a charge state of 2-4. Fragment mass tolerance was set to 20ppm. Methionine oxidation was set as variable modification, while cysteine carbamidomethylation was predefined by the software as fixed modification and could not be unselected even though the samples had been treated with methyl methanethiosulfonate (MMTS) to reversibly sulfenylate cysteine introducing beta-methylthiolation. However, since only around 8% of the peptides identified with MS Amanda 2.0 contained cysteines, this incorrect parameter was considered negligible in the course of this analysis, while using the correct fixed modification would have surely resulted in even better results for all CHIMERYS™ searches.

Peptide areas were quantified using IMP-apQuant^28^ using only PSM of high confidence level, with a minimum sequence length of 7 and a minimum score of 150 for MS Amanda 2.0 and -99 for the CHIMERYS™ Ion Coefficient and match between runs and RT correction were disabled. Retention time tolerance was set to 0.5, missing peaks to 2 and FWHM interpolation was enabled, and number of checked peaks set to 5. The results were filtered to 1% FDR on the protein level using the Percolator algorithm integrated in Thermo Proteome Discoverer. Statistical significance of differentially abundant peptides and proteins between different conditions was determined using a LIMMA test.^19^

## Supporting information

Supplemental Figure 1 - WWA optimization

Supplemental File 1 - Immunoprecipitation analyses

## ASSOCIATED CONTENT

### Supporting Information

Supporting information is available to download free of charge.

**Supplemental Figure 1**: Ideal precursor window size for wide window acquisition to optimally utilize CHIMERYS™ depends on sample abundance.

**Supplemental Table 1**: Chromatographic separation methods used

**Supplemental Table 2**: MS methods used

**Supplemental File 1**: Immunoprecipitation results

## AUTHOR INFORMATION

### Author Contributions

RM and MM conceptualized the study, performed experiments, data analysis and wrote the manuscript. RM and MM contributed equally to this work. AS prepared Smarca5 IP samples. GK and KS maintained, and reorganized LC-MS systems used for this work. FB conceptualized work on Smarca5. KM conceptualized the study and performed data analysis.

## ACKNOWLEDGMENT

This work supported by the EPIC-XS, Project Number 823839, funded by the Horizon 2020 Program of the European Union, by the project LS20-079 of the Vienna Science and Technology Fund and the by the ERA-CAPS I 3686, P35045-B, P32054 (FB) and P33380 (FB) project of the Austrian Science Fund. We thank the IMP for general funding and access to infrastructure and especially the technicians of the protein chemistry facility for continuous laboratory support. We are grateful to MSAID and Thermo Fisher Scientific in particular to Bernard Delanghe. and Martin Frejno for the opportunity to test CHIMERYS™ and Proteome Discoverer™ 3.0. We also thank Thermo Fisher Scientific for access to the micropillar array columns particularly Jeff Op de Beeck and Paul Jacobs.

