## Supplemental Figure 1 - WWA optimization for "Wide Window Acquisition and AI-based data analysis to reach deep proteome coverage for a wide sample range, including single cell proteomic inputs"

Supplemental Material

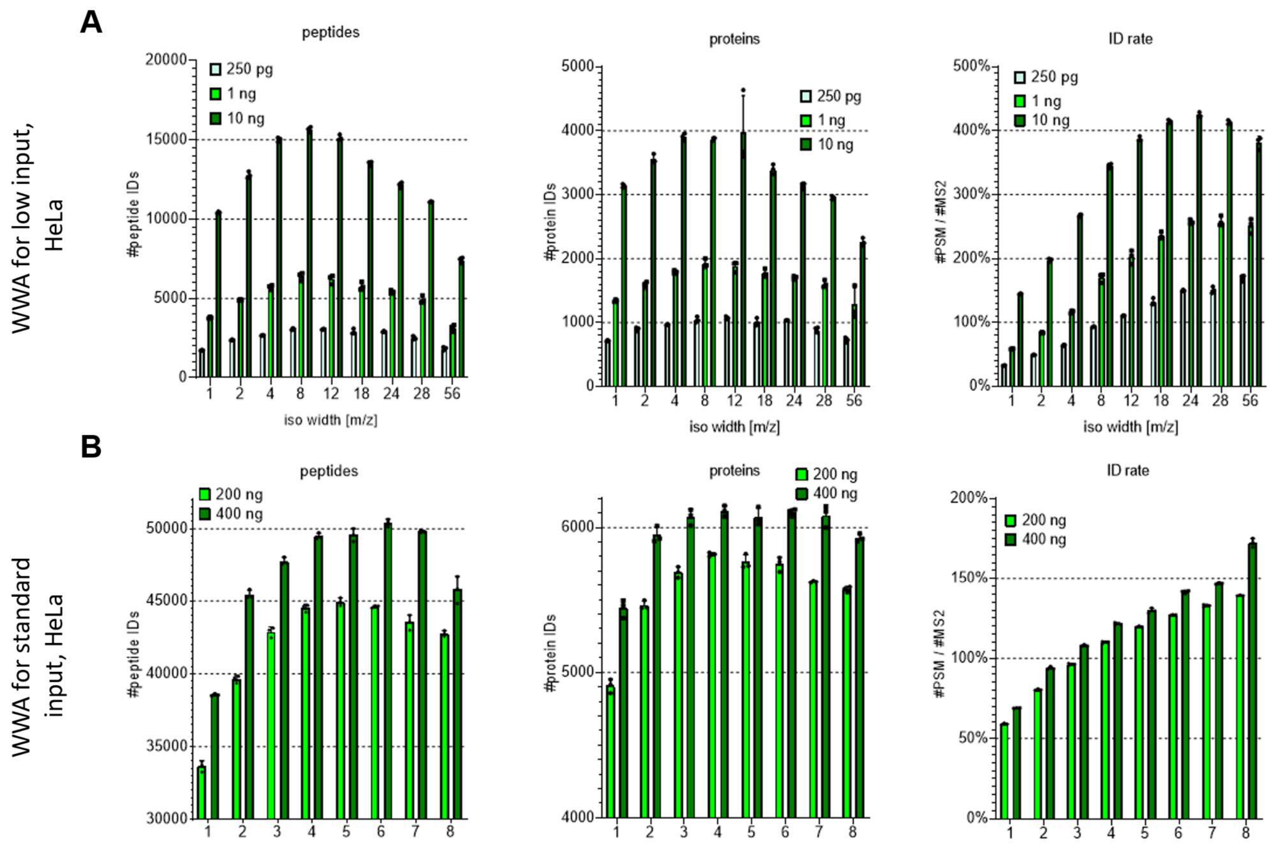


*Supplemental Figure 1:* ***Ideal precursor window size for wide window acquisition to optimally utilize CHIMERYS™ depends on sample abundance****. Different isolation window sizes were tested to identify the most well-suited precursor isolation window size for* ***(A)*** *low sample input such as 250 pg up to 10 ng measured on the 5.5 cm column and* ***(B)*** *standard sample input from 200 to 400 ng leading measured on the 50 cm column on peptide and protein level. ID rates were calculated by dividing the number of obtained peptide sequence matches (PSM) by the number of recorded MS2 spectra. ID rates over 100% indicate that on average more than one PSM could be identified per chimeric spectra. Bars indicate means and error bars indicate standard deviations, n= 3 technical replicates.*
